# Dose-controlled tDCS reduces electric field intensity variability at a cortical target site

**DOI:** 10.1101/793836

**Authors:** Carys Evans, Clarissa Bachmann, Jenny Lee, Evridiki Gregoriou, Nick Ward, Sven Bestmann

## Abstract

**Background:** Variable effects limit the efficacy of transcranial direct current stimulation (tDCS) as a research and therapeutic tool. Conventional application of a fixed-dose of tDCS does not account for inter-individual differences in anatomy (e.g. skull thickness), which varies the amount of current reaching the brain. Individualised dose-control may reduce the variable effects of tDCS by reducing variability in electric field intensities at a cortical target site.

**Objective:** To characterise the variability in electric field intensity at a cortical site (left primary motor cortex; M1) and throughout the brain for conventional fixed-dose tDCS, and individualised dose-controlled tDCS.

**Methods:** The intensity and distribution of the electric field during tDCS was estimated using Realistic Volumetric Approach to Simulate Transcranial Electric Stimulation (ROAST) in 50 individual brain scans taken from the Human Connectome Project, for fixed-dose tDCS (1mA & 2mA) and individualised dose-controlled tDCS targeting left M1.

**Results:** With a fixed-dose (1mA & 2mA), E-field intensity in left M1 varied by more than 100% across individuals, with substantial variation observed throughout the brain as well. Individualised dose-controlled ensured the same E-field intensity was delivered to left M1 in all individuals. Its variance in other regions of interest (right M1 and area underneath the electrodes) was comparable with fixed- and individualised-dose.

**Conclusions:** Individualized dose-control can eliminate the variance in electric field intensities at a cortical target site. Assuming that the current delivered to the brain directly determines its physiological and behavioural consequences, this approach may allow for reducing the known variability of tDCS effects.

## Introduction

Non-invasive electrical stimulation, including transcranial direct current stimulation (tDCS) uses a weak constant or alternating electrical current to influence neural activity in cortical networks [1,2]. This can modify behaviour [3–5] in both health and disease [6–9]. Unlocking the therapeutic potential however, is limited by the variable effects of stimulation, which could contribute to the small effect sizes often reported [10–13]. This known variability mandates novel approaches that increase the efficacy and reliability of the technique.

One major cause of variable effects and individual differences of tDCS are inter-individual variations in anatomy [14,15]. Anatomical variation, such as skull and cerebrospinal thickness, sulcal depth, and cortical folding, can influence the spread and intensity of current reaching the brain [16–19]. For example, skull conductivity strongly affects the amount and direction of current entering the brain [20–22].

Indeed, variation in E-field intensity at a cortical target region has been estimated to be around 100% [17]. Under the assumption that the E-field in a brain region directly relates to the physiological and behavioural effect of stimulation, this substantial variation alone could explain a large proportion of variable outcomes across individuals and studies.

Nevertheless, the conventional application of a fixed intensity of tDCS stimulator output (e.g. 1 or 2mA) across individuals ignores how much current actually reaches the brain in a given individual. This “one-size-fits-all” approach results in substantial variation in E-fields in cortex [20,23].

Reduction of inter-individual variability may be possible using current flow modelling [24–27]. This provides estimates of E-field across an individual’s brain, given a specific electrode montage and stimulator output. Recent studies using intracranial recordings in animals and human patients demonstrate a close qualitative correspondence between model estimates and measured E-fields [28–32]. These models may therefore provide the opportunity to quantify how much current is delivered into the brain, and where.

However, few studies have utilised current flow modelling to control the dose of the E-field actually delivered within and across individuals. Most commonly, these models have been used for individualising the electrode montage to target a specific cortical target [24,30]. For example, Dmochowski and colleagues [28] found a 64% increase in electric current in a cortical target and 38% improvement in a behavioural task using individualised tDCS montages in stroke patients. However, with this method of dose-control the intensity of E-field in the target region is not fixed across individuals.

Individualising electrode montage, such as in stroke patients [28] also substantially increases variability in the spatial distribution of current flow across brain regions and individuals. Here we made the assumption that the physiological and behavioural effects of tDCS primarily arise from a specifically targeted cortical region.

Here we propose a simple and alternative method of dose-control, where stimulator output is individualised and electrode montage is fixed across individuals. Using current flow models, one can determine the tDCS intensity needed to deliver a target E-field intensity to a given cortical target site in each individual (see Figure 1 for a visualisation of this concept). Whilst this approach does not eliminate variance in current flow distribution, it may result in more consistent distribution of current than if variable electrode montages were used.

**Figure 1.**
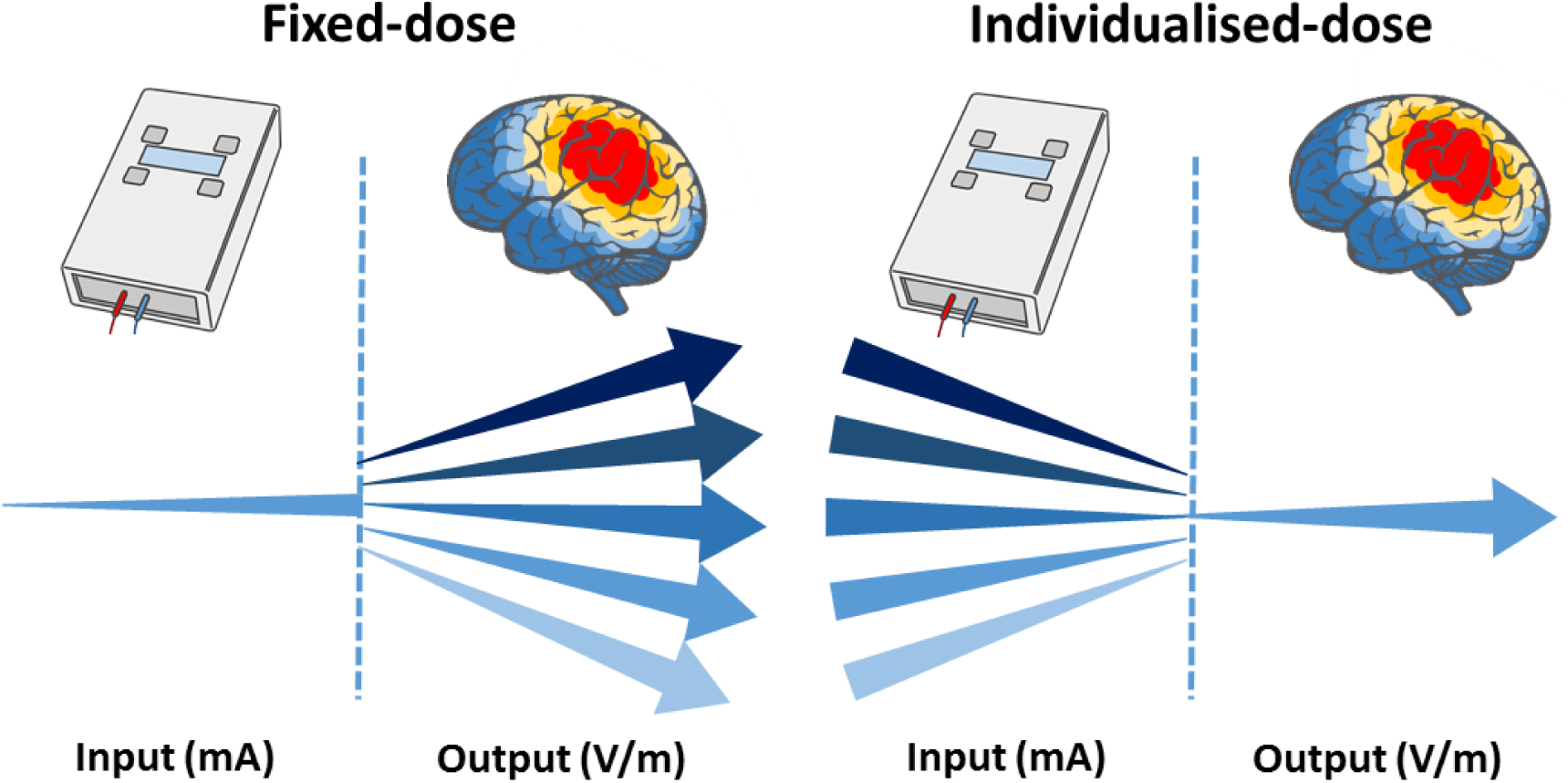
A fixed-dose of tES results in variable E-field distribution across the brain and intensity delivered to the cortical target site (e.g. left M1). Individualising-dose results in variable stimulator output for each individual, but constant electrical field intensity at the cortical target site.

The present study formally quantified and compared E-field intensity and distribution across the brain when delivering (i) a fixed intensity of tDCS to all individuals, and (ii) dose-controlled tDCS delivering the same E-field at a cortical target site across individuals. Specifically, we applied tDCS over the left primary motor cortex, one of the most common targets of electrical stimulation [1,33–35]. These models were conducted using a ‘directional’ electrode montage, in which the electrodes are placed anteriorly and posteriorly to the primary motor cortex (M1) such that current flow is directed perpendicularly to the central sulcus. This montage has been used to minimize variation in the direction of current flow at the target site [36–38]. We compared this with the ‘conventional’ electrode montage used to target M1, in which the anode is placed over M1 and the cathode over the contralateral forehead [1,39].

We confirm that delivery of conventional fixed-dose stimulation varies E-field intensity at a cortical target site by more than 100% between two groups. These intensities can vary by a factor of two with relatively small sample sizes (N=15). Moreover, this approach leads to substantial variation in E-field distribution across the brain. Critically, E-fields at intensities thought sufficient for physiological targeting are widespread and consistently encompass the regions between stimulation electrodes.

By contrast, dose-controlled tDCS ensures the same E-field intensity is delivered to a cortical target site, here left M1. However, the spread of current remains widespread. We argue that reducing the variable outcomes of non-invasive electrical stimulation requires individualised dose-control or vastly increased sample sizes [40,41].

## Materials and Methods

### Structural MRIs

T1-weighted structural MRIs from 50 randomly selected healthy adults (aged 22-35, 23 males, 27 females) were obtained from the Human Connectome Project (HCP) database (https://ida.loni.usc.edu/login.jsp). Images were acquired from a 3.0 T Siemens Connectome Skyra scanner using a standard 32 channel Siemens receiver head coil. Scanning parameters were as follows: spatial resolution: 0.7mm isotropic voxels, repetition time (TR): 2400ms, TE: 2.14ms, TI: 1000ms, flip angle: 8 degrees, field of view: 224 × 224mm using Siemens AutoAlign feature, iPAT: 2. Head movements were tracked using an optical motion tracking system (Moire Phase Tracker, Kenticor)[42].

The HCP project is supported by the National Institute of Dental and Craniofacial Research (NIDCR), the National Institute of Mental Health (NIMH) and the National Institute of Neurological Disorders and Stroke (NINDS). HCP is the result of efforts of co-investigators from the University of Southern California, Martinos Center for Biomedical Imaging at Massachusetts General Hospital (MGH), Washington University, and the University of Minnesota.

### Current Flow Modelling

We estimated current flow throughout cortex in individual brains using Realistic Volumetric Approach to Simulate Transcranial Electric Stimulation (ROAST) software package, version 2.7 (https://www.parralab.org/roast/) [26]. ROAST is an end-to-end pipeline that automatically processes individual MRI volumes to generate a 3D rendering of E-field distribution based on user-defined tDCS protocol. ROAST accepts a 1mm^3^ resolution MRI image, which is segmented using SPM12 (https://www.fil.ion.ucl.ac.uk/spm/) into grey matter, white matter, cerebrospinal fluid (CSF), bone, skin, and air cavities. Automatic touch-up of the segmented images to remove holes is completed using morphological operations and simple heuristics (detailed in [26,30]). Electrodes are then placed on the scalp surface based on user-defined 10-10 coordinates. Using iso2mesh [43], a volumetric mesh is generated from 3D multi-domain images to generate the finite element model. The model is then solved for voltage and E-field distribution using getDP [44]. Conductivity values for each tissue type are as follows (in S/m): grey matter: 0.276; white matter: 0.126; CSF: 1.65; bone: 0.01; skin: 0.465; air: 2.5 × 10^−14^; gel: 0.3; electrode: 5.9×10^7^.

In order to extract E-field intensity and distribution in the brain, structural MRI and E-field images produced by ROAST were pre-processed using SPM12. Structural and E-field images were normalised (resampled to 2×2×2mm voxels) into standard space (Montreal Neurological Institute; MNI template) and then smoothed using a 4mm full-width at half-maximum Gaussian kernel. Images were smoothed to increase signal to noise ratio and improve accuracy of observed E-fields when conducting group-level analyses. An explicit binary mask of grey and white matter was created using the normalised structural images in order to constrict subsequent analyses to voxels within the brain. First, average grey and white matter tissue masks were created using the non-binary tissue masks produced by ROAST for each individual during segmentation. These masks were then transformed into binary grey and white matter tissue masks, with an inclusion threshold of >0.1 intensity. Binary grey and white matter masks were then combined to create one mask of the brain. See Figure 2 for complete pipeline. Finally, an anatomical template was generated to visualise the data by averaging the structural MRI images of all individuals.

**Figure 2.**
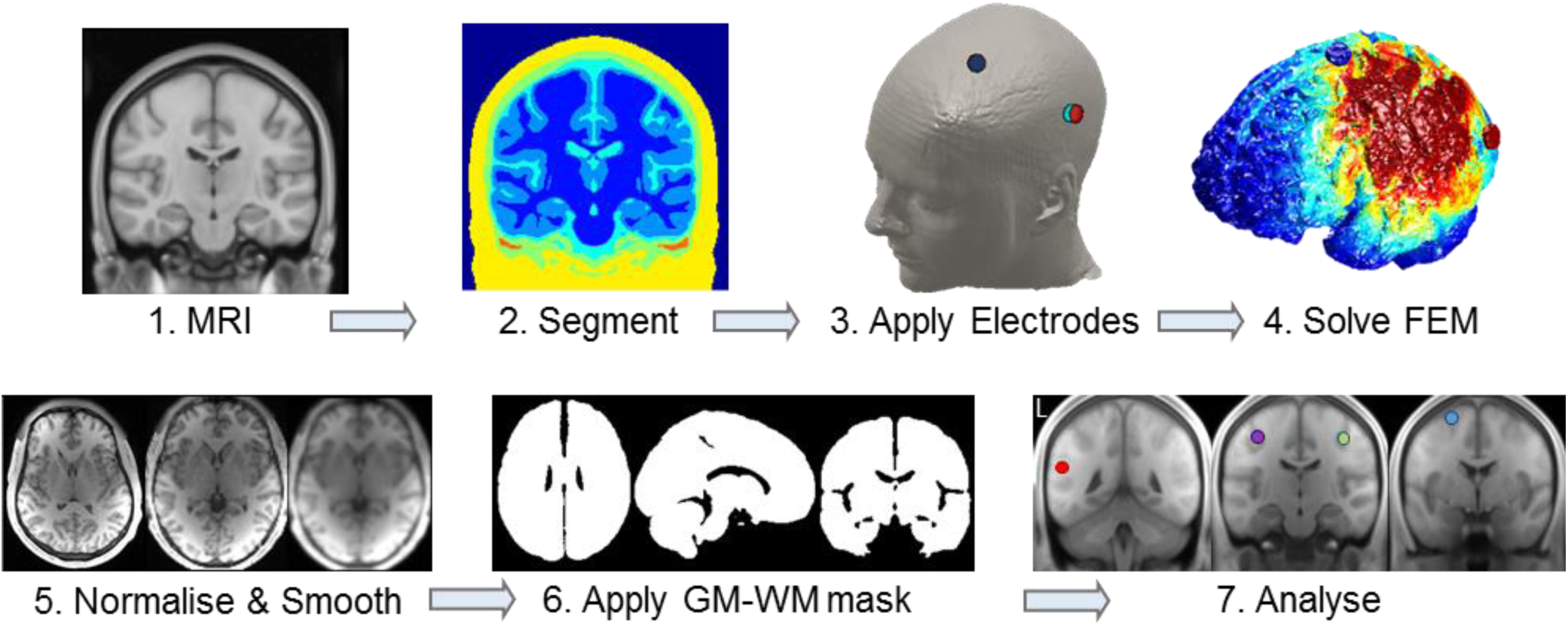
Current flow modelling pipeline. Steps 1-4 are automated in ROASTv2.7 to generate the E-field distribution: based on a structural MRI image (1), each tissue type is segmented (2), electrodes are applied (3), and the finite element model (FEM) is solved (4). Using SPM12, the structural MRI and E-field images are then normalised and smoothed (5), a grey and white matter explicit mask is created (6), and applied to the E-field to constrict analyses to within the brain (7). Step 7 shows the regions of interest: left M1 (purple), right M1 (green), and cortical sites under the electrodes: left angular gyrus underneath CP5 (red) and left premotor cortex underneath FC1 (blue).

### tDCS protocol

Current flow models simulated bipolar application of transcranial direct current stimulation (tDCS) using disc electrodes (6mm radius, 2mm height). Electrodes were placed over 10-10 coordinates CP5 (anode) and FC1 (cathode). This ‘directional’ electrode montage was selected to target the hand region of the left primary motor cortex (M1), by placing electrodes in a posterior-anterior orientation perpendicular to, and either side of M1 [36,45].

#### Fixed-dose

across two models, the amplitude of stimulator output, and hence the modelled dose were fixed at 1mA and 2mA. These intensities are commonly used in tDCS applications and are known to induce acute excitability changes [46,47]. Average intensities of electric current (V/m) were then extracted from left M1 across the sample, and used to determine the desired target intensities in left M1.

#### Individualised-dose

the sample averages for 1mA (M=0.185 ± 0.033V/m) and 2mA (M=0.369 ± 0.064V/m) target intensities in left M1 were 0.185V/m and 0.369V/m respectively. We set these values as target intensities in left M1 for the dose-controlled simulations (i.e. to reduce variance in E-field intensity at the cortical target site). For these current flow models we individualised stimulator output (Individualised_dose_) such that the targeted intensity occurred in left M1 for each individual:

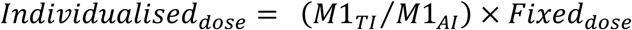

Where M1_TI_ is the Target Intensity of E-field in left M1 across individuals, M1_AI_ is the Actual Intensity when applying a fixed-dose (1mA/2mA) for each individual, and Fixed_dose_ is the fixed stimulator output that corresponds to the individualised-dose (e.g. target intensity based on the average intensity when 1mA fixed-dose is applied). The calculated individualised-dose was rounded to the nearest 0.025mA to adhere to parameter limitations of the constant current stimulator produced by NeuroConn (DC-Stimulator Plus; NeuroConn, Ilmenau, Germany).

### Data Analysis

Statistical analyses on current flow models produced by ROAST were carried out using MATLAB (The MathWorks, Inc., Natick, MA, USA) and SPM12. Alpha level was 0.05 and analyses were corrected for multiple comparisons using family-wise error correction (FWE).

#### Regions of interest (ROI)

to assess variance in E-field intensity between individuals in target left M1 (MNI: -38, -20, 50), eigenvariates were extracted from regions of interest (5mm radius). MNI coordinates for left M1 were based on previous activation likelihood estimations [48]. Additional ROIs included the cortex underneath electrode sites, the angular gyrus (AnG; MNI: -54, -46, 22), underneath CP5 and premotor cortex underneath FC1 (PMC; MNI: -24, 4, 66), and right M1 (MNI: 30, -18, 48; [48]). ROIs for CP5 and FC1 were converted from Talairach coordinates taken from anatomical locations of 10-10 projections [49], and then extended deeper within the cortex to avoid inclusion of CSF. MNI coordinates specify the centre of each ROI (see Figure 2 for ROIs).

#### Reproducibility of left M1 intensities with typical sample-sizes

based on the average E-field intensity observed in left M1 when 1mA fixed-dose was applied, the likelihood of smaller samples yielding similar intensities was investigated. 1000 bootstrap samples of 15 and 30 individuals (i.e. 45,000 resamples in total) were analysed, reflecting sample sizes typically observed in tDCS studies [40,50]. Each iteration of resampling randomly drew individuals from the original sample (N=50) with replacement. For each bootstrap sample, the average intensity in left M1 was obtained.

#### Spatial distribution of E-field

one-sample t-tests of E-field images were conducted to identify brain regions where the E-field was significantly above zero. The distribution is described qualitatively, as manipulating the variance in individualised-but not fixed-dose models meant the data was no longer statistically comparable. Topography was explored for the whole sample and two example individuals.

#### Probability maps of E-field intensities above 0.14V/m

to determine the distribution of E-field intensities above 0.14V/m throughout the brain in a large population, we generated probability maps for 1mA fixed-dose and the corresponding individualised-dose (0.185V/m target intensity in M1). For the individualised-dose models, the E-field images for each subject were summed, and the lowest E-field intensity within the left M1 ROI of the summed image was selected (0.14V/m). For both the fixed- and individualised-dose models, we then created binary images from each individual’s normalized E-field images, which included voxels >0.14V/m. We then created probability maps of E-field values exceeding 0.14V/m, by summing these binary images. The summed values were converted into percentage to create the final probability maps.

#### Comparing E-fields for directional and conventional electrode montages

E-field intensities within ROIs and the spatial distribution of E-fields were measured for the ‘conventional’ electrode montage [1,39]. These were qualitatively compared to the directional electrode montage. Conventional montage models were conducted for 1mA fixed-dose and individualised-dose to obtain 0.182V/m in left M1 (based on sample averages for 1mA: M=0.182 ± 0.036). ROIs included target left M1, right M1, and cortex under the electrodes: the left post central gyrus (PCG; MNI: -44, -22, 50) underneath C3, and right anterior prefrontal cortex (aPFC; MNI: 24, 52, 14) underneath FP2.

## Results

### Dose-controlled tDCS reduces variance in E-field intensity in left M1

For the directional electrode montage (CP5-FC1) and fixed stimulator output, E-field intensities were highly variable across individuals. Intensities ranged from 0.125-0.249V/m with 1mA (M=0.185V/m ± 0.033), and 0.263-0.518V/m with 2mA (M=0.369V/m ± 0.064). E-field intensities in left M1 therefore varied by ∼100% with a given stimulator output.

Interestingly, the most consistent intensities in each model were observed in the right hemisphere (i.e., contralateral to stimulation). The global maximum (i.e. the statistical maximum with least variance in E-field) for 1mA was in the right medial temporal lobe (MNI: 38, -12, -16, *t*_(1,49)_=82.70, *p*<.001). For 2mA, the global maximum was located in parahippocampal gyrus (MNI: 28, 2, -30, *t*_(1,49)_=83.47, *p*<.001). However, while consistent, the E-field intensity in these regions was very low for both 1mA (M=0.060V/m ± 0.005) and 2mA (M=0.097V/m ± 0.009).

Dose-control vastly removed variance in E-field intensity in left M1. Intensities ranged from 0.176-0.190V/m with 0.185V/m target intensity (M=0.184V/m ± 0.003), and 0.353-0.386V/m with 0.369V/m (M=0.368V/m ± 0.006). However, this variance was due to rounding the individualised-dose to the nearest 0.25mA and repeating the ROAST pipeline. Taking this into account, these results suggest that variance in E-field intensities at the target location was essentially eliminated. See Figure 3 for intensities in left M1 and Table 1 for descriptive statistics.

**Table 1.**
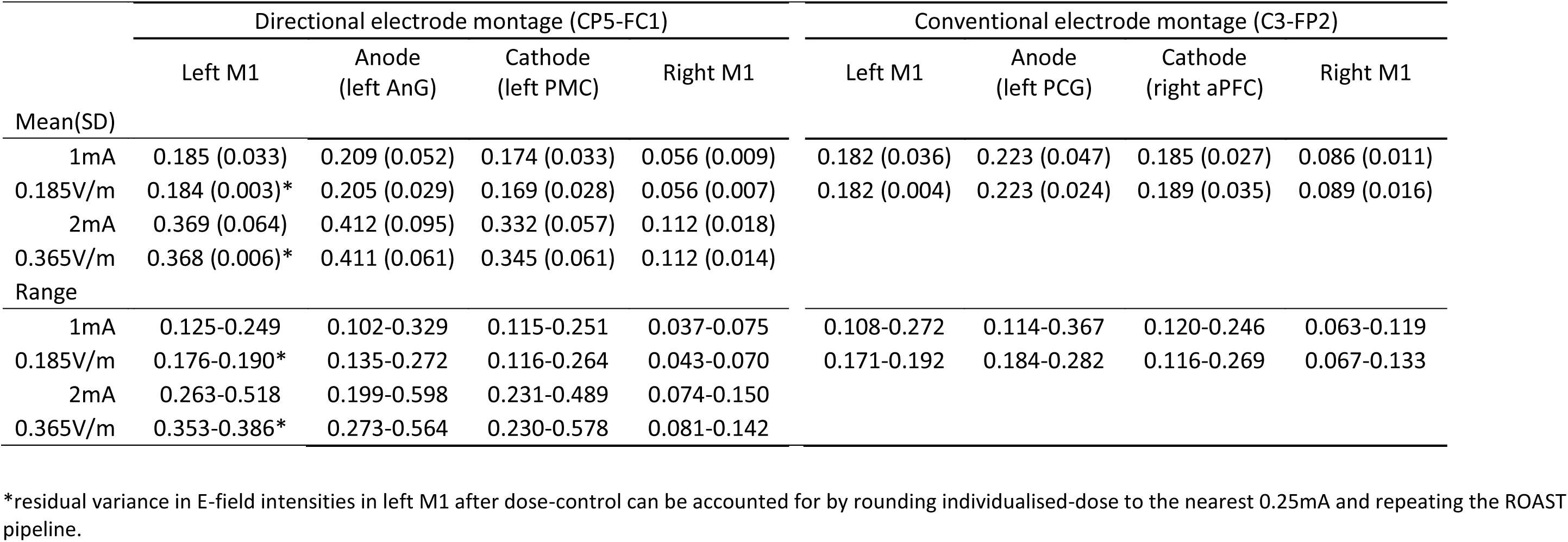
E-field intensities (V/m) in target left M1 and additional ROIs after a fixed-dose (1mA & 2mA) and individualised-dose to obtain 0.185V/m and 0.365V/m.

|  | Directional electrode montage (CP5-FC1) |  |  |  | Conventional electrode montage (C3-FP2) |  |  |  |
| --- | --- | --- | --- | --- | --- | --- | --- | --- |
|  | Left M1 | Anode<br>(left AnG) | Cathode<br>(left PMC) | Right M1 | Left M1 | Anode<br>(left PCG) | Cathode<br>(right aPFC) | Right M1 |
| Mean(SD) |  |  |  |  |  |  |  |  |
| 1mA | 0.185 (0.033) | 0.209 (0.052) | 0.174 (0.033) | 0.056 (0.009) | 0.182 (0.036) | 0.223 (0.047) | 0.185 (0.027) | 0.086 (0.011) |
| 0.185V/m | 0.184 (0.003)* | 0.205 (0.029) | 0.169 (0.028) | 0.056 (0.007) | 0.182 (0.004) | 0.223 (0.024) | 0.189 (0.035) | 0.089 (0.016) |
| 2mA | 0.369 (0.064) | 0.412 (0.095) | 0.332 (0.057) | 0.112 (0.018) |  |  |  |  |
| 0.365V/m | 0.368 (0.006)* | 0.411 (0.061) | 0.345 (0.061) | 0.112 (0.014) |  |  |  |  |
| Range |  |  |  |  |  |  |  |  |
| 1mA | 0.125-0.249 | 0.102-0.329 | 0.115-0.251 | 0.037-0.075 | 0.108-0.272 | 0.114-0.367 | 0.120-0.246 | 0.063-0.119 |
| 0.185V/m | 0.176-0.190* | 0.135-0.272 | 0.116-0.264 | 0.043-0.070 | 0.171-0.192 | 0.184-0.282 | 0.116-0.269 | 0.067-0.133 |
| 2mA | 0.263-0.518 | 0.199-0.598 | 0.231-0.489 | 0.074-0.150 |  |  |  |  |
| 0.365V/m | 0.353-0.386* | 0.273-0.564 | 0.230-0.578 | 0.081-0.142 |  |  |  |  |

\*residual variance in E-field intensities in left M1 after dose-control can be accounted for by rounding individualised-dose to the nearest 0.25mA and repeating the ROAST pipeline.

**Figure 3.**
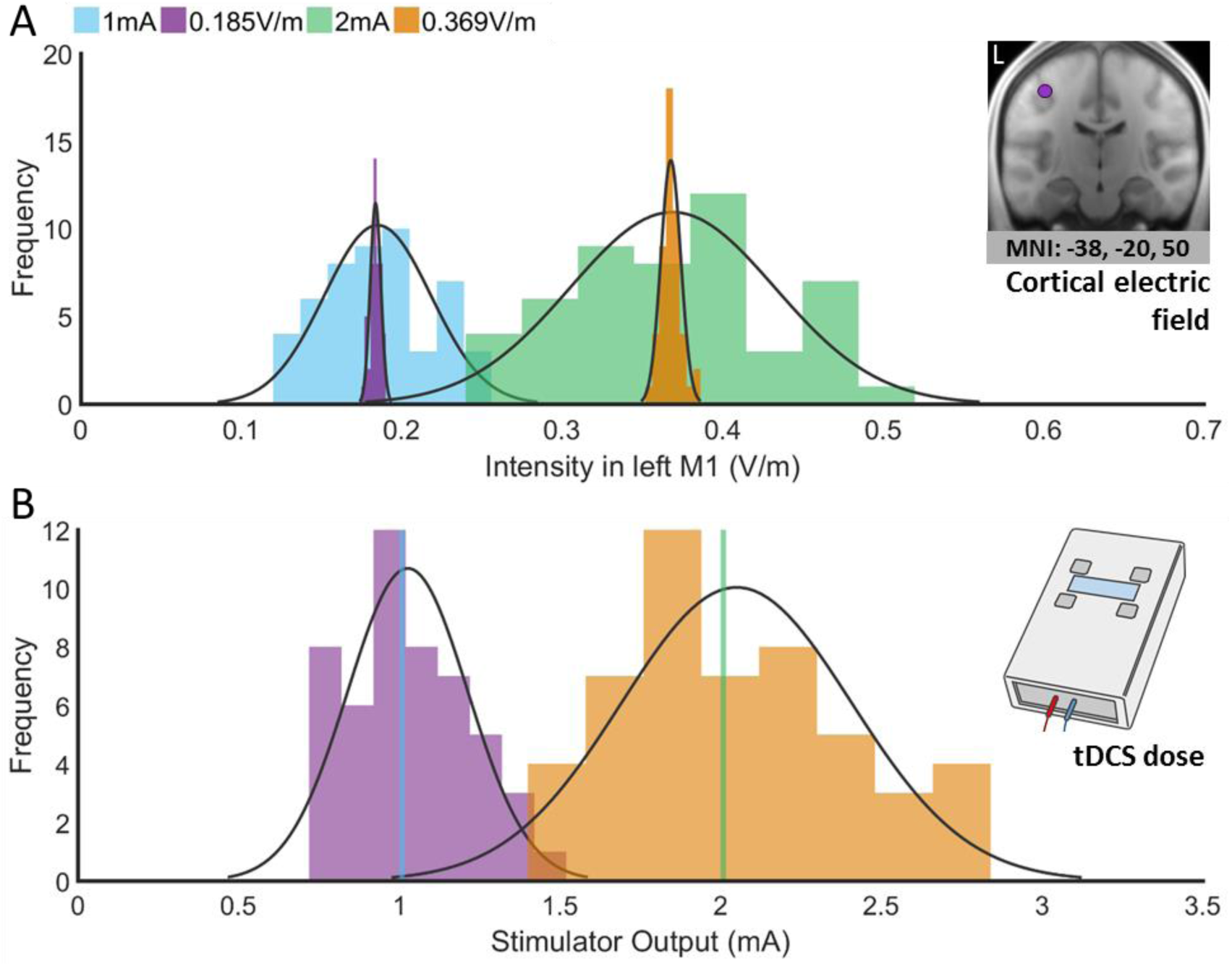
(A) Distribution of E-field intensity (V/m) in left M1. **(B)** Stimulator output (mA). Data shown for 50 individuals when applying a fixed-dose of 1mA (blue) and 2mA (green), and individualised-dose to obtain 0.185V/m (purple) and 0.369V/m (orange) in left M1. Applied to each model are fitted normal probability density functions. Coronal section of the brain marks ROI (MNI: -38, -20, 50) for left M1 (purple).

Consequently, the least variance in E-field was detected around the left motor cortex/precentral gyrus (MNI: -36, -20, 46) for both 0.185V/m (*t*_(1,49)_=114.52, *p*<.001) and 0.369V/m (*t*_(1,49)_=111.43, *p*<.001), and was comparable to target intensities 0.185V/m (M=0.174 ± 0.009) and 0.369V/m (M=0.348 ± 0.019). By individualising-dose, the E-field was therefore most consistent near the target site, as expected and as per design, and reached the specified target intensities for this region.

This means that stimulator output differed substantially across individuals (Figure 3). To ensure delivery of 0.185V/m to left M1, stimulator output ranged from 0.750-1.475mA (M=1.027 ± 0.187), and ranged from 1.425-2.8mA (M=2.050mA ± 0.358) for a target of 0.369V/m.

### Poor consistency of left M1 E-field intensities with fixed-dose and small sample-sizes

Bootstrapped samples of 1mA fixed-dose produced wide sample-to-sample variability in left M1 E-field intensity. For small samples (*N=*15), E-field variance ranged between 28-100% within each resample, and 52% of 1000 resamples yield a mean <0.185V/m. For larger samples (*N=*30), E-field variance within each resample ranged from 58-100%, and 48% of 1000 resamples produced a mean <0.185V/m.

In the sample with the lowest mean, 80% of intensities in left M1 were <0.185V/m (M=0.158V/m ± 0.031, N=15; M=0.167V/m ± 0.033, N=30), whilst in the samples with the highest mean, the majority of individuals had intensities higher than 0.185V/m (M=0.213V/m ± 0.031, N=15; M=0.202V/m ± 0.33, N=30) of intensities were <0.185V/m (see Figure 4). Therefore, when applying tDCS with a fixed output intensity, the mean intensity of current delivered to a cortical target site can substantially vary across different samples, and this variation in turn is exacerbated for small sample sizes.

**Figure 4.**
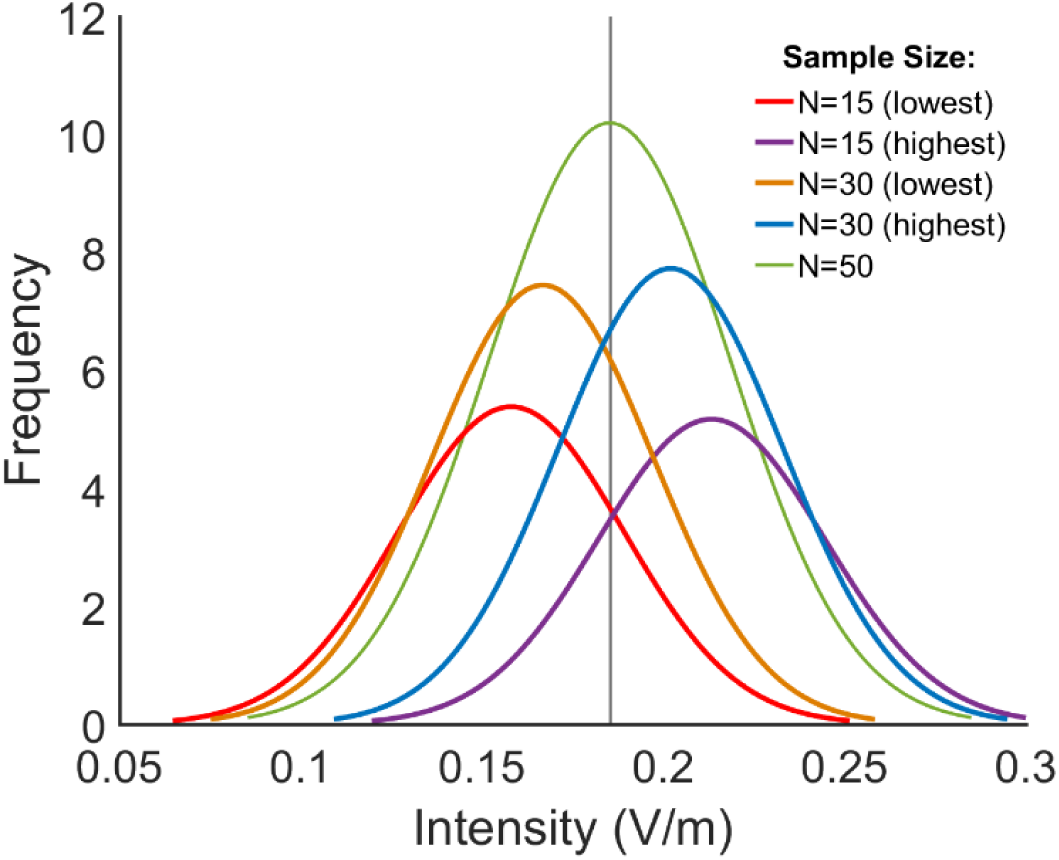
Example distributions of E-field intensity in left M1 with 1mA fixed-dose for different sample sizes. The original sample (N=50; green) used in this study is compared to bootstrapped samples with the lowest and highest mean E-field intensities for sample sizes of N=15 and N=30, respectively. The vertical line indicates average intensity in the original sample after 1mA: 0.185V/m. This illustrates that for small sample sizes, the mean intensities at a cortical target site can vary up to 100% across groups of individuals.

### Dose-control does not reduce variance in E-field intensity in non-target regions

Having explored variance in E-field intensity in the target site left M1, the variation in E-fields across the brain was assessed. We focussed on the left AnG (underneath CP5), PMC (underneath FC1), and right M1.

#### Regions under stimulation electrodes (left AnG and left PMC)

Fixed-dose yielded highly variable E-fields in left AnG and PMC. In left AnG, intensities ranged from 0.102-0.329V/m with 1mA, and 0.199-0.598V/m with 2mA. In left PMC, intensities ranged from 0.115-0.251V/m with 1mA, and from 0.231-0.489V/m with 2mA. Notably, these intensities exceeded those observed in left M1.

Individualised-dose showed a comparable range and intensity of E-field. With 0.185V/m target intensity, intensities in left AnG ranged from 0.135-0.272V/m, and from 0.273-0.564V/m with 0.369V/m. In left PMC, intensities ranged from 0.116-0.264V /m with 0.185V/m, and from 0.230-0.578V/m with 0.369V/m. While dose-controlled tDCS delivery, as done here, removes E-field variance in the target region, it does not therefore greatly alter E-field variance underneath the stimulation electrodes or elsewhere (Figure 5, Table 1).

**Figure 5.**
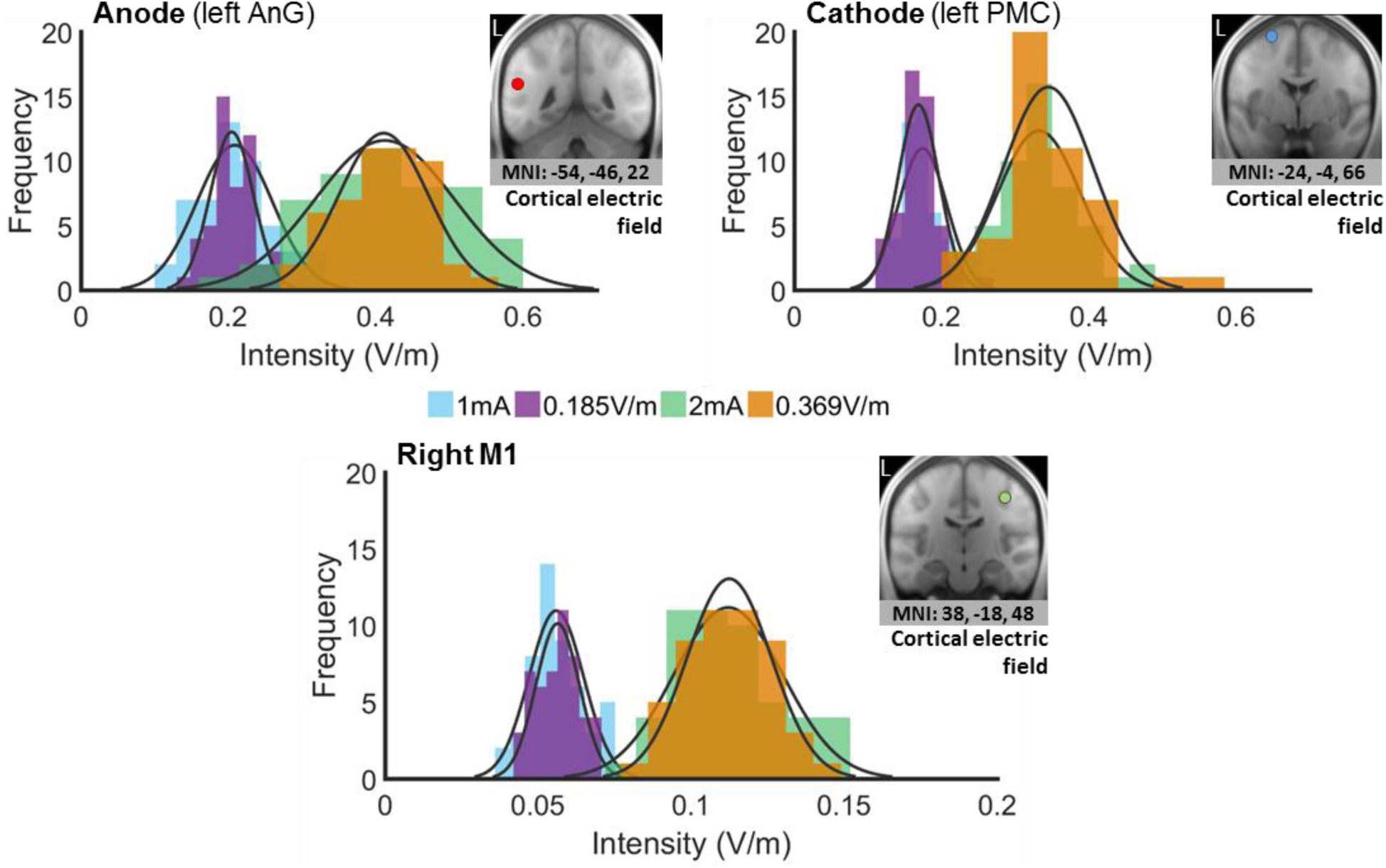
Distribution of E-field intensity (V/m) at each ROI (left AnG, left PMC, right M1) across 50 individuals. Intensities are shown for fixed-dose 1mA (blue) and 2mA (green) and individualised-dose to obtain 0.185V/m (purple) and 0.369V/m target intensity (orange) in left M1. Fitted normal probability density functions are displayed for each model. Note: right M1 x-axis covers a smaller range.

#### Right M1

1mA fixed-dose produced low E-field intensities in contralateral M1, ranging from 0.037-0.075V/m. When increasing stimulator output to 2mA, as is now commonly done for M1 stimulation [51,52], E-fields reached intensities similar to those observed in targeted left M1 after 1mA stimulation (range: 0.074-0.150V/m).

Similarly, for dose-controlled tDCS, intensities in right M1 were low for a left M1 target intensity of 0.185V/m (range: 0.043-0.070V/m), but reached intensities between 0.081-0.142V/m for a target intensity in left M1 of 0.369V/m. At high stimulation intensities, both fixed- and individualised-dose application of tDCS can therefore result in substantial contralateral E-field intensities.

### Diffuse but more consistent spatial distribution of E-field in the brain with dose-control

One-sample t-tests confirmed that tDCS induces a large and complex topography of E-field above zero, including both cortical and subcortical structures.

Applying 1mA fixed-dose, intensities >0.1V/m extended from parietal to frontal and temporal regions across individuals. E-fields were predominantly restricted to the left hemisphere, with some extension into right hemisphere white matter structures via the corpus callosum. At 2mA, the E-field extended further into inferior regions of the left hemisphere including cerebellum and brainstem. Intensities >0.1V/m were observed in contralateral frontoparietal grey matter, contralateral white matter, and subcortical structures (Figure 6).

**Figure 6.**
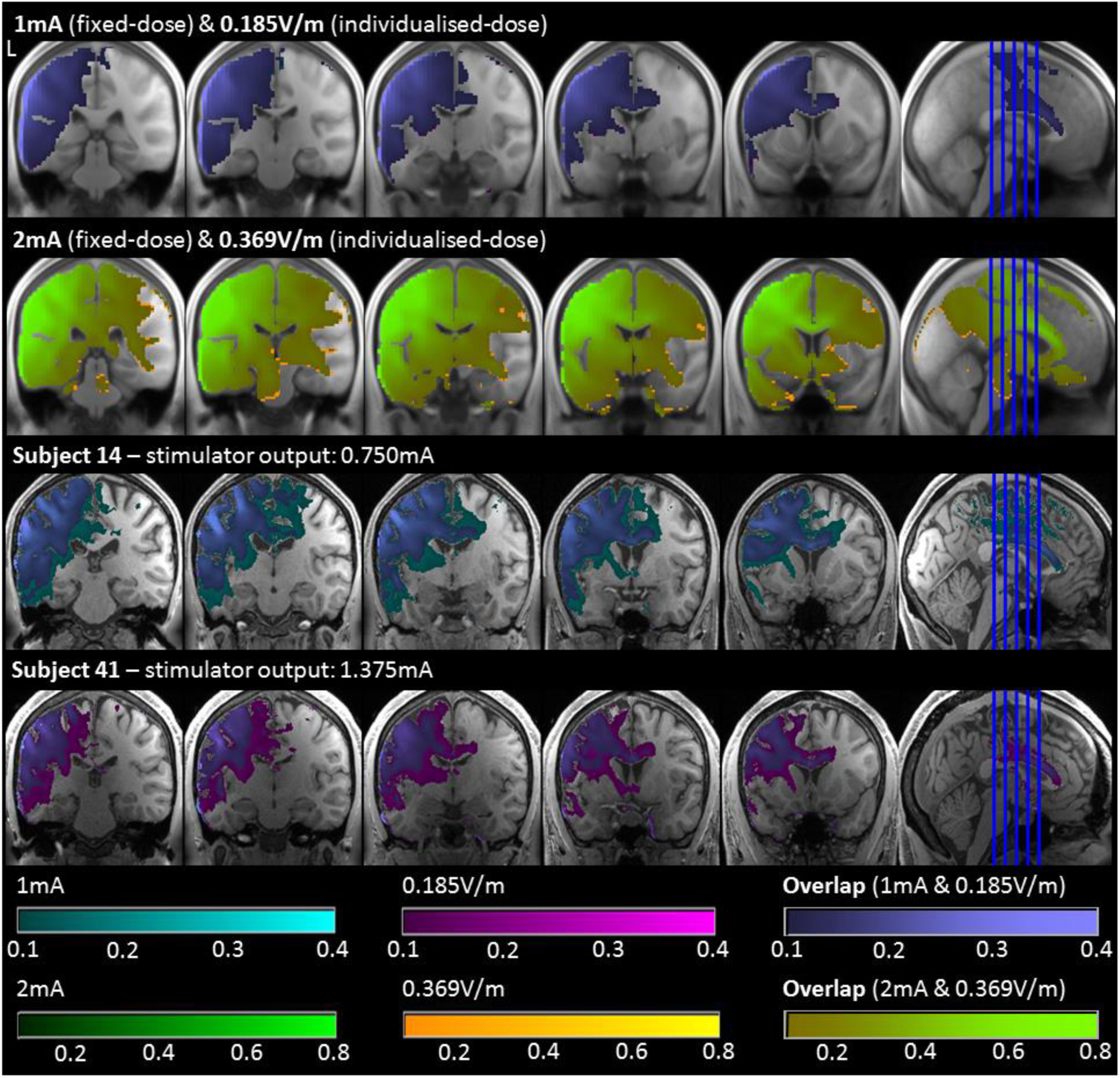
E-fields (V/m) of 50 individuals and subject 14 and 41. Shown are 1mA fixed-dose (cyan) and individualised-dose to obtain 0.185V/m (pink) with E-field overlap in purple (image threshold: 0.1-0.4), and 2mA fixed-dose (green) and individualised-dose to obtain 0.369V/m (yellow) with overlap represented in lime (image threshold: 0.1-0.8).

Similar distributions of E-fields were observed after dose-controlled tDCS, and did not extend to additional cortical areas.

E-field distribution in individuals exhibiting high (e.g., subject 14) or low (e.g., subject 41) intensities in left M1 after 1mA fixed-dose illustrate the heterogeneous distribution of E-fields across individuals. The E-field in subject 14 extended medially, including much of the left hemisphere’s grey and white matter, and left temporal regions, whereas the E-field in subject 41 was less diffuse, with intensities >0.1V/m largely contained in left parietal grey matter (see Figure 6).

However, with dose-control, the E-field distribution was comparable in these subjects, matching the distribution observed across the whole sample. This illustrates that E-field distribution becomes more consistent when individualising dose.

Our probability maps corroborate this observation. By combining binary masks of E-field >0.14V/m for each individual, these maps determined the likelihood at each voxel that E-fields would exceed this threshold across a large sample. Figure 7 shows that for both fixed- and individualised-dose, intensities >0.14V/m encompassed much of the lateral surface between electrodes in the left hemisphere in over 80% of individuals. However, individualising-dose considerably increased the number of regions where intensities exceeded 0.14V/m in 100% of individuals. The number of individuals with intensities >0.14V/m in temporal and contralateral structures also reduced with dose-control. Most notably, in target left M1, the number of individuals with intensities >0.14V/m was highly variable when applying a fixed-dose. With individualised-dose, intensities >0.14V/m within the ROI increased to approximately 90-100% of individuals, confirming that dose-control also reduces variability in a cortical target site.

**Figure 7.**
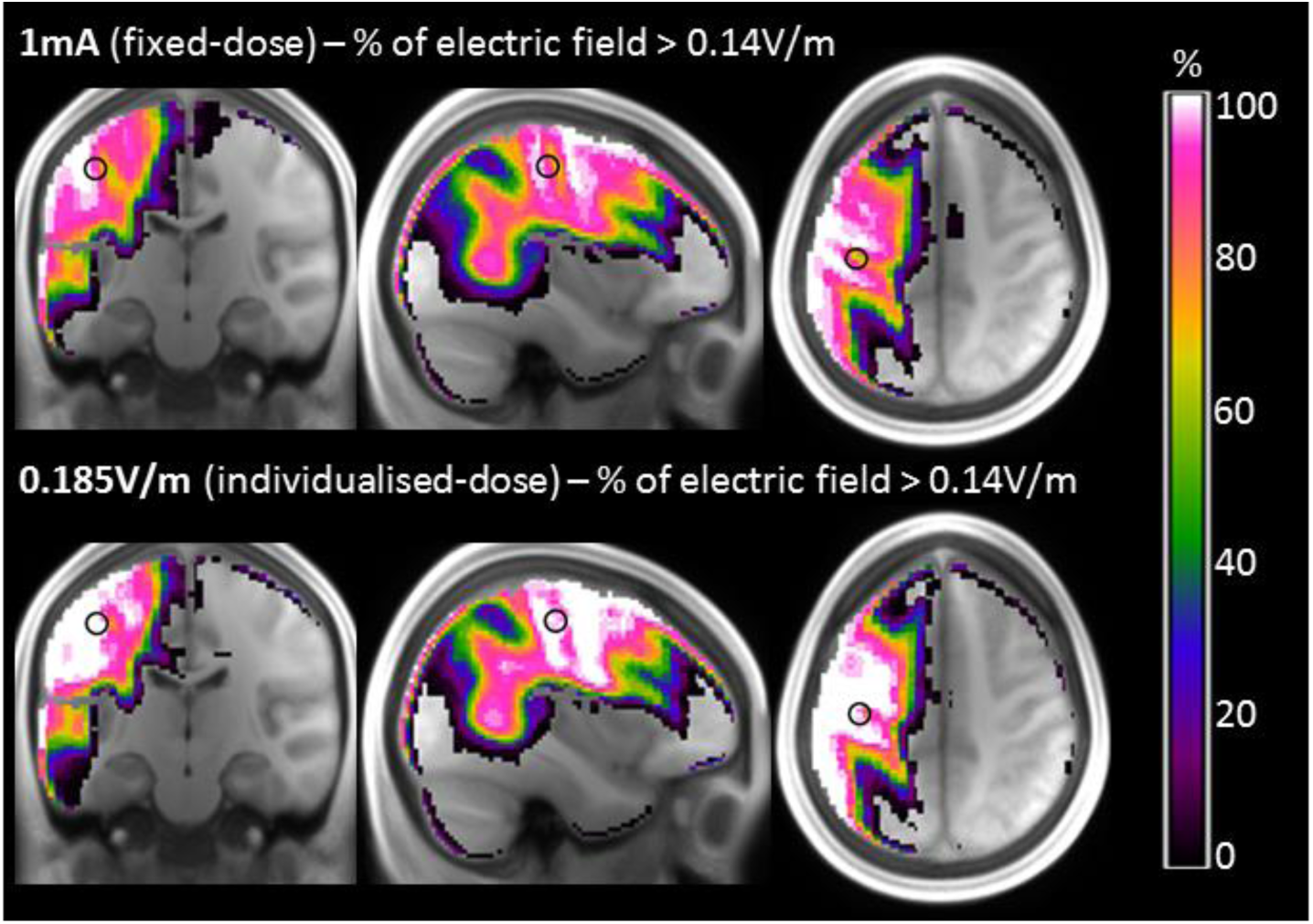
Probability maps of electric field exceeding 0.14V/m in 50 individuals when applying 1mA fixed-dose and individualised-dose to obtain 0.185V/m in left M1. Circled is the ROI for target left M1 (MNI: -38, -20, 50). Maps were generated by adding binary masks of electric field >0.14V/m for each individual.

### Conventional (C3-FP2) and directional (CP5-FC1) montages produce similar E-field intensities in target left M1 and ROIs

Applying 1mA fixed-dose with the conventional electrode montage (C3-FP2), intensities in left M1 were similar to those observed with the directional montage (CP5-FC1), ranging from 0.108-0.272V/m (M=0.182V/m ± 0.036). The global maximum for 1mA indicated that the most consistent E-field intensities were observed in anterior left cerebral white matter (MNI: -12, 18, 42, *t*_(1,49)_=79.37, *p*<.001) and were of comparable magnitude to left M1 (M=0.172V/m ± 0.015; range: 0.130-0.202V/m).

The reduction in variance of E-field intensity when using dose-control was comparable between conventional and directional montages. With a target intensity of 0.182V/m, intensities in left M1 ranged from 0.171-0.192V/m across individuals (M=0.182V/m ± 0.004). The least statistical variance in E-field was also in left M1 after dose-control (MNI: -38, -20, 50, *t*_(1,49)_=161.78, *p*<.001). For controlling dose in M1, stimulator output intensities ranged from 0.675-1.700mA (M=1.041mA ± 0.220). This demonstrates that for controlling the E-field delivered to a cortical target site, a fixed stimulator output is not a viable approach (Figure 8).

**Figure 8.**
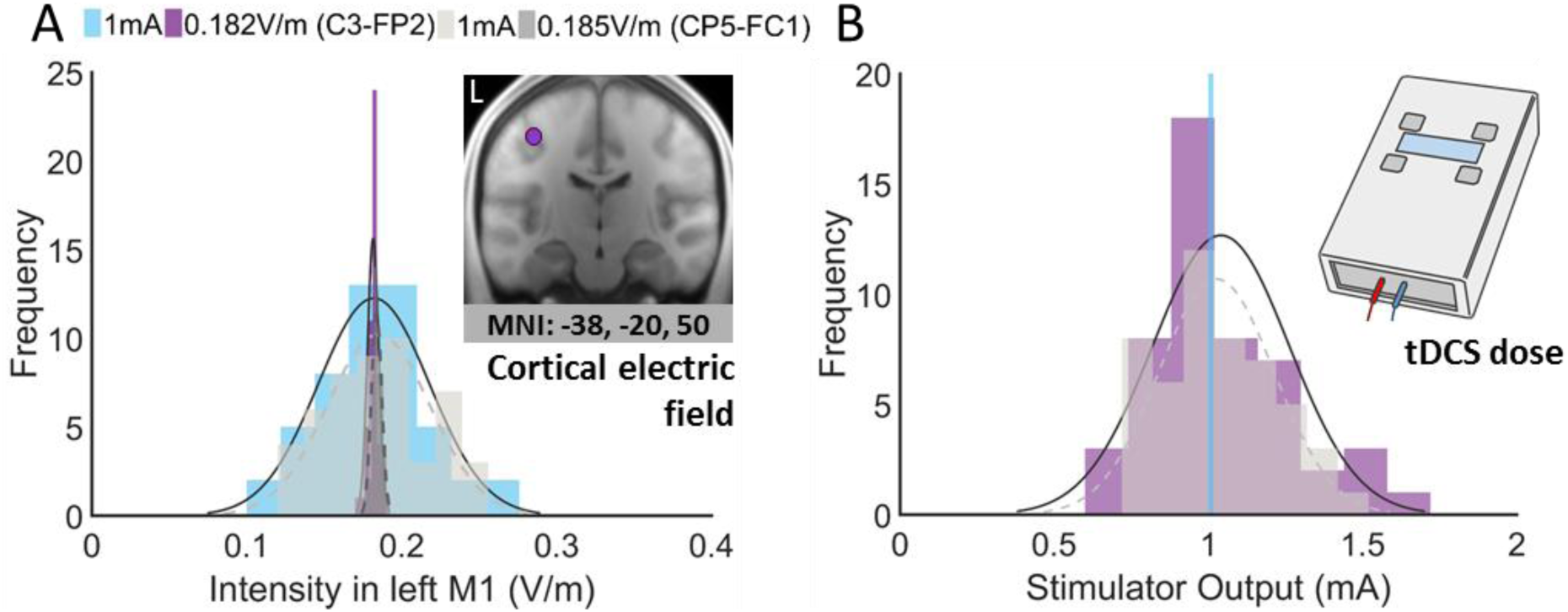
**(A)** Distribution of E-field intensity (V/m) in left M1 using conventional electrode montage (C3-FP2). Overlaid in grey are the intensity distributions for the directional montage (CP5-FC1). **(B)** Stimulator output (mA). Data shown for 50 individuals when applying a fixed-dose of 1mA (blue) and individualised-dose to obtain 0.182V/m (purple) in left M1. Applied to each model are fitted normal probability density functions; grey dashed lines denote normal distributions for directional montage models. Coronal section of the brain marks ROI (MNI: -38, -20, 50) for left M1 (purple).

#### Regions under stimulation electrodes (left PCG and right aPFC)

in line with the directional montage, fixed-dose tDCS produced highly variable E-fields in left PCG and right aPFC. Range and intensity of E-field in these regions was maintained when applying an individualised-dose to obtain 0.182V/m target intensity in left M1 (see Table 1).

#### Right M1

in contralateral M1, E-field intensities were lower, but importantly still greater than those observed in models using the directional montage when applying either a fixed- or an individualised-dose. Notably, there is overlap between the highest intensities observed in contralateral (right) M1 and the lowest intensities observed in left M1. To the extent that one assumes stimulation affects the targeted M1 region, this suggests that in a subgroup of individuals, stimulation can also influence M1 contralateral to the targeted hemisphere (see Figure X).

### Conventional (C3-FP2) electrode montage leads to greater spatial distribution of E-field in the brain

Figure 9 shows that E-field intensities exceeding 0.1V/m encompass much of the targeted (left) hemisphere when applying 1mA fixed-dose using a conventional M1 electrode montage. By contrast, for the directional montage, these intensities are largely restricted to the left hemisphere, suggesting that even though intensities in the targeted M1 region are comparable, there are considerable differences in the spatial extent of E-fields across the brain, including subcortical structures.

**Figure 9.**
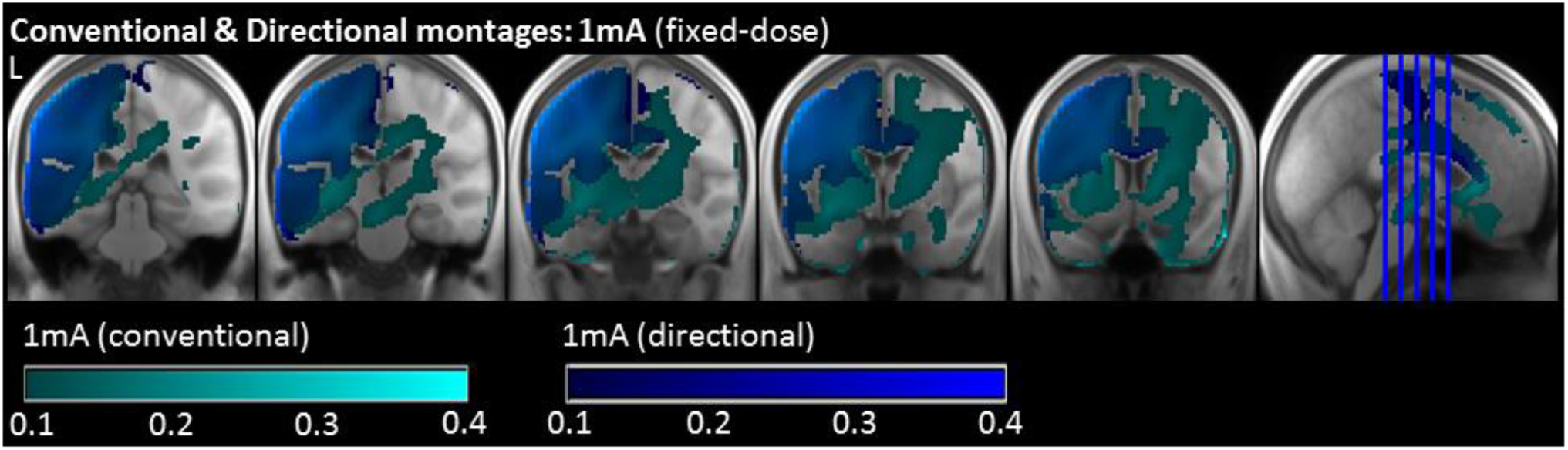
E-fields (V/m) of 50 individuals when applying 1mA fixed-dose using the conventional electrode montage (C3-FP2; cyan) and directional montage (CP5-FC1; blue). Image threshold: 0.1-0.4.

### Summary

Current flow models confirmed that fixed-dose tDCS yields substantially variable E-field intensities in left M1, and that the mean intensity across groups can vary by a factor of two with smaller sample sizes. Individualised dose-control can eliminate this variance. Yet, intensities in other ROIs are comparable across fixed- and individualised-dose models.

E-field distribution in the brain is widespread, extending contralaterally with higher stimulation doses or when applying lower doses using the conventional electrode montage. Heterogeneous distribution across individuals with fixed-dose tDCS can be greatly reduced when individualising-dose.

## Discussion

A central concern for widespread application of tDCS in health and disease is the known variability of physiological and behavioural outcomes, both between individuals and research studies [10–13]. Given the assumption that current entering the brain relates directly to the effect of stimulation, variance in E-field will be a major source of variability in tDCS effects. Here, we quantified that inter-individual differences in anatomy can lead to variation in E-field by as much as 100%. However, this variance can be controlled by individualising the dose of tDCS to deliver a fixed cortical E-field.

### Dose-control reduces variance in E-field intensities at a cortical target site

When applying a fixed-dose of tDCS, E-field intensities are heterogeneous both across different regions of the brain within an individual, and also across individual subjects. At the cortical target site (left M1), intensities varied by over 100%, which is consistent with previous estimates of E-field variability [17]. Typical application of a fixed-dose of tDCS is therefore likely to lead to variable physiological or behavioural effects across individuals [16–20,51].

With smaller sample sizes (∼15-30 individuals) commonly used in basic and translational tDCS studies [40,50,53], this heterogeneity can result in cohorts with entirely different stimulation profiles. Bootstrapped samples illustrated that, on average, for smaller samples the overall intensities in left M1 can vary substantially from our larger population mean. Therefore, it is not unreasonable to assume that the effectively applied current across studies may vary by a factor of two or more.

Assuming a linear relationship between current delivered to a cortical target and its physiological and behavioural consequences, this variation likely contributes to variable outcomes and hinders reproducibility across studies. Strikingly however, despite much work on modelling current flow induced by electrical stimulation, formal quantification of E-field variance is rarely reported (notable exceptions being [17,22,54]). Controlling for this source of variability must therefore be a priority.

Here we developed a simple and accessible pipeline using current flow modelling [26] and group-level statistics to quantify E-field in the brain when dose-controlling tDCS. Based on intensities in a cortical target (left M1) observed in our cohort, tDCS dose was individualised to produce a fixed cortical E-field in this region across all individuals. In other words, rather than applying the usual fixed stimulator output, this output is adjusted for each individual to deliver the desired target intensity at a cortical target site. Consequently, stimulator output varied by 100% across individuals, but stayed within the safe range (≤4mA) for tDCS applications in a healthy population [46].

Notably, dose-control did not reduce variance or spread of E-field in other regions. This is entirely expected given the linear properties of Ohm’s law. Treating head tissues as pure resistors means that, for example, maintaining electrode montage while simulating a twofold increase in injected current will result in an identical distribution of E-field across the brain, but with a twofold increase in intensity at each cortical location. Specifically, in cortical areas underneath the stimulation electrodes, and in contralateral (right) M1, intensities varied by ∼100% or more for both fixed- and individualised-dose models.

Interestingly, higher doses of tDCS (∼2mA) resulted in intensities in contralateral right M1 approaching similar amplitudes to those observed in left M1 when lower doses were applied (1mA). Higher tDCS intensities may target distal regions that are not underneath or in-between the electrodes, including homotopic contralateral areas. Likewise, when using the conventional electrode montage (C3-FP2) there was overlap in the highest intensities observed in contralateral (right) M1 and the lowest intensities in target left M1, suggesting that stimulation can influence both target and contralateral M1 in a subgroup of individuals. This observation seems particularly relevant for the recent debate about the possible non-linear effects of tDCS intensity on physiology and behaviour [51,55,56]. For example, if increasing tDCS intensity effectively alters contralateral M1 excitability in a subset of individuals, M1-M1 interactions may contribute to the seemingly non-linear (or non-monotonic) changes observed in the targeted M1 excitability.

### Dose-control reduces variance in spatial distribution of E-field across the whole brain

As expected, our results confirmed that both fixed- and individualised-dose produce comparable E-field distributions across the brain, given a fixed electrode montage. With lower doses (∼1mA), intensities over 0.1V/m encompassed large proportions of the left hemisphere, whereas higher doses (∼2mA) also included contralateral regions. However, using dose-control these distributions become qualitatively more comparable.

These observations corroborate previous reports that tDCS produces diffuse E-field in the brain [18,24,27,57]. Validation studies reported peak intensities of 0.5V/m [32] and 0.4V/m [31] with 1mA transcranial electrical stimulation, and 0.8V/m when applying 2mA tDCS [31]. These intensities correspond to those observed at the cortical surface in the present study. We note that irrespective of whether these values are quantitatively accurate, our results remain qualitatively valid: fixed-dose tDCS results in large E-field variance at a cortical target site and across the brain, which is removed by individualising dose.

In the present study, we applied dose-control with the assumption that physiological or behavioural effects of tDCS relate to E-field intensities delivered to a cortical target in a straightforward (linear) way. In other words, the E-field in a brain region of interest directly determines the likelihood of physiological and behavioural consequences, whether this is an increase or decrease in excitability. We therefore here controlled E-field delivered to this target region.

However, other parameters such as current flow direction, spatial extent, or gradient of polarisation across the cortical surface [58] may be relevant. For example, other brain regions receiving intensities above a certain threshold will likely contribute to the effect of stimulation. In particular, probability maps demonstrate that across the sample, a wide range of cortical regions between electrode sites consistently reach E-field intensities thought to be sufficient for physiological targeting. Future work will address how these parameters could be controlled for.

Individualised application of tDCS likely also requires consideration of differences in structural or functional state of the brain, particularly in ageing [59,60] or clinical populations such as stroke [25,61,62]. We would argue that such development should precede optimization of protocols based on the number of stimulation sessions [63]), intensity [56,64], or individual differences including baseline physiological [65,66] or cognitive function, such as performance ability [67] or attention [68].

Our approach uses a straightforward calculation of the required stimulator output given a desired target intensity in a cortical target site. This is distinct from other optimisation approaches determining optimum electrode placement for maximal intensity in a cortical target [18,29,69]. Yet, as the effects of tDCS likely arise from interaction of larger networks given the non-focal distribution of current, these may vary substantially for different electrode montages. Further, while numerical optimisation has been proposed to achieve this [70], it is not commonly available. Moreover, we establish a pipeline for potentially calculating normative datasets for E-field distribution throughout the brain, given a specific electrode montage. In principle this can be achieved with any number of individual brains in a straightforward way, which is relevant given the cost and difficulty obtaining individual brain scans required for current flow modelling. Normative distributions in large cohorts also enable required sample sizes to be formally derived.

Going forward, the utility of current flow modelling requires further investigation, namely to determine whether these models can indeed be used to reduce the physiological or behavioural variability associated with conventional tDCS delivery. As previously noted [20], there is a surprising lack of such validation in the field, which will inevitably determine the efficacy of current flow modelling. Given that intensity of stimulation influences the degree of membrane polarisation [13,51,55], our prediction is that controlling the delivery of current to the brain will reduce variance in the effects of tDCS.

Recent studies suggest that current flow in a cortical target can predict response to transcranial electrical stimulation [71,72]. For example, Laakso and colleagues [71] proposed that variability in motor evoked potentials could be partly explained by E-fields in M1 induced by tDCS. Specifically, these authors pointed at the putative relationship between the normal component of E-field in the hand area of M1. As ROAST uses volume rather than surface based segmentation to produce current flow models, it may be beneficial in future work to compare the explanatory power of both volumetric and boundary element methods when determining the effects of tDCS.

Application of our dose-control approach to other electrode montages, such as high definition (HD)-tDCS [29,73], or other target sites, will in the future provide a framework for validation of other ways to optimise current delivery.

### Conclusion

The present study demonstrates the substantial variability in E-field reaching the brain due to inter-individual differences in anatomy. It also provides a straightforward approach to reduce variance by dose-controlling tDCS using current flow modelling.

This effectively shifts variance from E-field in the cortex to stimulator output. If dose-control also reduces variance in stimulation effects, the efficacy of non-invasive electrical stimulation, including tDCS could be increased.

## Acknowledgements

Structural MRI data for this project was provided by the MGH-USC Human Connectome Project (HCP; Principal Investigators: Bruce Rosen, M.D., Ph.D., Arthur W. Toga, Ph.D., Van J. Weeden, MD). HCP funding was provided by the National Institute of Dental and Craniofacial Research (NIDCR), the National Institute of Mental Health (NIMH), and the National Institute of Neurological Disorders and Stroke (NINDS). HCP data are disseminated by the Laboratory of Neuro Imaging at the University of Southern California

